# Automated cellular structure extraction in biological images with applications to calcium imaging data

**DOI:** 10.1101/313981

**Authors:** Gal Mishne, Ronald R. Coifman, Maria Lavzin, Jackie Schiller

## Abstract

Recent advances in experimental methods in neuroscience enable measuring in-vivo activity of large populations of neurons at cellular level resolution. To leverage the full potential of these complex datasets and analyze the dynamics of individual neurons, it is essential to extract high-resolution regions of interest, while addressing demixing of overlapping spatial components and denoising of the temporal signal of each neuron. In this paper, we propose a data-driven solution to these challenges, by representing the spatiotemporal volume as a graph in the image plane. Based on the spectral embedding of this graph calculated across trials, we propose a new clustering method, Local Selective Spectral Clustering, capable of handling overlapping clusters and disregarding clutter. We also present a new nonlinear mapping which recovers the structural map of the neurons and dendrites, and global video denoising. We demonstrate our approach on in-vivo calcium imaging of neurons and apical dendrites, automatically extracting complex structures in the image domain, and denoising and demixing their time-traces.

## 1. Introduction

Detecting numerous small regions of interest is a prevalent problem in biomedical imaging applications, where images are composed of hundreds of structures such as cells, organelles or neurons, acquired in various imaging sensors such as imaging mass cytometry (Cytof), Multiplexed ion beam imaging (MIBI), hyperspectral microscopy, etc. [1, 2]. In these datasets each pixel in the image is associated with a high-dimensional vector of measurements and a critical task is the ability to extract small structures from these datasets. This challenge also exists in biomedical videos, such as calcium imaging data, where each pixel is associated with a high-dimensional time-trace of up to tens of thousands of time-frames.

Calcium imaging, based on high resolution genetically encoded calcium indicators combined with two photon microscopy, has revolutionized the ability to track the activity of neuronal networks in awake behaving mammals. These methods not only enable chronic, minimally invasive recordings of large populations of neurons with cellular level resolution, they also allow recordings from identified neuronal subtypes [3]. This ability has major significance, and is expected to raise understanding of neuronal circuits to a new level, since different neuronal populations receive and send specific inputs associated with distinct network functions.

However, the analysis of this data relies first and foremost on the ability to automatically extract high-resolution regions of interest (ROI) of each neuron and their corresponding time-traces. Given these traces it is then possible to recover the spiking activity of each neuron from the slower dynamics of the calcium indicator for further analysis [4–6]. One challenge of ROI extraction is the separation of overlapping ROIs, where the overlap is due to the projection of a 3D volume onto a 2D imaging plane. A second challenge is the images themselves, which suffer both from varying dynamic range, and low signal-to-noise ratio due in part to the activation of calcium in the neuropil creating a noisy heterogeneous background [7]. In addition the data suffers from measurement noise and movement artifacts [8].

Analysis of calcium imaging data has typically relied on the assumption that ROIs are spatially sparse (occupying a small region of pixels in the image plane), with a sparse temporal signal. Some methods address only the spatial extraction of the ROIs, without recovering the temporal signal of each ROI. In [9], ROIs are detected by filtering a local cross-correlation peak image computed from the temporal correlation of each pixel with its adjacent neighbors. Pachitariu et al. [7] develop a generative spatial model for biological images based on convolutional sparse block coding. Applied to calcium imaging, they learn models of somas and dendrites from a single image of the mean temporal activity.

Other methods address both the spatial and temporal components, identifying both spatial ROIs and their time-traces. Mukamel et al. [10] proposed spatiotemporal independent component analysis (ICA), using skewness as a measure of sparsity. Diego and Hamprecht [11] extend convolutional sparse coding to video data, extracting sparse spatial components and their sparse temporal activity while estimating a non-uniform and temporally varying background. More recent approaches focus on matrix factorization with different constraints and penalties on the background component, sparseness of the solution and temporal dynamics [12–15]. Pnevmatikakis et al. [14] presented a state of the art method based on constrained matrix factorization, which explicitly models the calcium indicator dynamics. More recent work has deviated from sparsity-based solutions towards general model-free and data-driven approaches [8,16], and our framework detailed in this paper falls within this line of work. Our extraction does not rely on modeling the fluorescent calcium activity but takes a general approach to solving clustering of high-dimensional data in the presence of noisy clutter. It should be noted that despite all these algorithmic advances, manual segmentation is still commonplace, requiring expensive expert time and yielding unreproducible data.

In this paper, we develop a graph-based approach for high-dimensional clustering, motivated from a manifold learning perspective. Our main contribution in this paper is three-fold. First, we propose a new approach, Local Selective Spectral Clustering (LSSC), that differs from the popular algorithm and its variations [17–22], by both relying on localized viewpoints in the embedding space, and by looking deeper into the spectrum. Second, we obtain a structural map of the neurons, demonstrating a depth-map recovery of the data. We demonstrate that this map is more informative and cleaner than the typically used correlation image. Finally, we develop a full framework for ROI extraction and denoising, without imposing sparsity constraints. Our framework results in ROI extraction of neuronal structures, a temporal denoising of the time-traces for each ROI and as a by-product, enables spatial denoising which provides an enhanced viewing of the video itself. In contrast to other methods, we make no assumptions on the morphology of the ROIs or the statistics of the temporal signal.

Our approach can be applied to both long-running experiments and to datasets following a fixed-length trial protocol. Such data is especially appropriate as the repetitive nature of the trials leads to both stronger spatial and temporal connections. We demonstrate our approach on multi-trial experiments of fixed-length in awake mice, and demonstrate the ability to identify, extract and demix neurons and apical dendrites from dense fluorescent images.

This paper is organized as follows. We introduce relevant notations and briefly review nonlinear spectral embeddings in Sec. 2. In Sec. 3 we present the LSSC approach, and its application to calcium imaging. Sec. 4 introduces a local denoising scheme for the ROIs and a global denoising scheme for the video itself. In Sec. 5 we present results on in-vivo neuronal recordings of somas and dendrites.

## 2. Background

Our approach is general and can be applied to a wide range of high-dimensional biomedical and remote sensing imagery. Here we focus on calcium imaging where the high-dimensionality arises from the temporal nature of the measurements. While we derive our notations in the spatiotemporal context, our approach can be generalized for multi-channel imagery, such as hyperspectral images. The notations in this paper follow these conventions: matrices **M** are denoted by bold uppercase and vectors **v** are denoted by bold lowercase.

Let *Z*(*x, y, t, n*) be a spatiotemporal dynamical system where *x, y* are the spatial coordinates, *t* : 0 ≤ *t* ≤ *T* is a short time scale representing a trial of fixed length *T* and *n* is an index of the separate trials. Denote by *Z_n_*(*x, y, t*) the measurement at pixel location (*x, y*) and time *t* for the *n*-th trial. The matrix 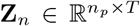 is the flattening of the 3D tensor of a single trial *n* into a matrix whose rows are indexed by *i* ∈ {1, …, *n_p_*}, and whose columns are indexed by the time-frames in the trial, where *n_p_* is the total number of pixels in the image plane. Each row in the matrix is the time-series of pixel *i*, or location (*x_i_, y_i_*), denoted by **z***_n,i_* = **z**_*n*_(*x_i_, y_i_*) = (*Z_n_*(*x_i_, y_i_,* 1), …, *Z_n_*(*x_i_, y_i_, T*))*^T^* ∈ ℝ*^T^*. Specifically, we focus on calcium imaging data, where *Z* are fluorescent measurements.

Our goal is to find spatial ROIs, denoted {*C_k_*}, which are subset of pixels in the image plane that compose a neuronal structure, such that ∀*i, j* ∈ *C_k_*, **z***_n,i_* and **z***_n,j_* are similar for some *n*. We assume that there may be overlapping ROIs due to the projection of the 3D volume to the 2D imaging plane, so that the ROIs are not disjoint, i.e., there may ∃*k, k^’^* such that *C_k_* ∩ *C_k_’* ≠ ∅. We require that an ROI *C_k_* is a connected component in the image plane.

Having identified spatial ROIs, the temporal measurements of each pixel in an ROI *C_k_* are used to calculate a time-trace of its fluorescent activity, denoted *F_k_*(*t, n*). These time-traces serve further analysis of the identified ROIs, for example, spike extraction [4–6], or uncovering dynamical factors in the neuronal activity [23, 24]. To this end, the time-traces of the neurons need to be demixed (for overlapping neurons) and denoised.

### 2.1. Nonlinear embedding

We aim to both extract distinct ROIs and denoise the data. Spectral methods have been used successfully for both of these tasks, such as spectral clustering and diffusion maps, [18,25,26]. Specifically, we will focus on the eigenvectors of a spatial random-walk affinity matrix on the data. We calculate a global embedding, which we will employ both for non-local denoising of the video [27] and a spatial clustering for ROI extraction.

Let each pixel *i* in the image plane be a node in a graph for which we observe a high-dimensional feature vector: the time series **z**_*n,i*_. To define the weights between nodes, we disregard spatial proximity but rather focus on similarity of the temporal measurements. For each trial *n*, we define the affinity matrix **K***_n_* whose elements are the pairwise affinity between the *T*-length time-series of two spatial locations **z***_n,i_* and **z**_*n,j*_:

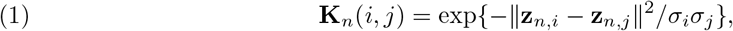

where *σ_i_, σ_j_* are the local self-tuning scale [19] of **z***_n,i_* and **z***_n,j_*, respectively. Due to the large size of the dataset, *n_p_* pixels, we calculate **K***_n_* as a sparse symmetric affinity matrix, using *k*-nearest neighbors [28].

A corresponding row-stochastic matrix is obtained by normalizing the rows of **K**_*n*_:

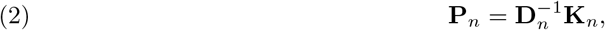

where **D***_n_* is a diagonal matrix whose diagonal elements are the degree of each node: **D***_n_*[*i, i*] = Σ*_j_* **K***_n_*[*i, j*]. The matrix **P***_n_* can be considered the transition matrix of a Markov chain on the data, where **P***_n_*[*i, j*] is the probability of jumping from **z***_n,i_* to **z***_n,j_*. The eigen-decomposition of **P***_n_* yields a a sequence of eigenvalues 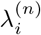 such that 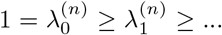, and bi-orthogonal left and right eigenvectors, 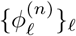 and 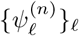, respectively.

We calculate the eigenvectors of **P***_n_* for each trial *n* ∈{1, …, *N* }, obtaining *N* different nonlinear embeddings of the spatial graph, denoted by Ψ*_n_*: ℝ *^T^* →ℝ*^d^*. Each pixel is associated with its corresponding elements in the set of right eigenvectors:

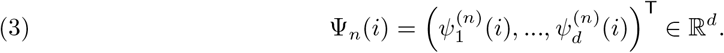

Thus, each pixel is no longer represented by a high-dimensional vector of temporal activity, but rather the values in the embdding space. Note that unlike diffusion maps [26], we do not weight the eigenvectors by their corresponding eigenvalues. As opposed to principal component analysis (PCA), which is commonly used in neuroscience and has been used for ROI extraction [10], the representation here is nonlinear and derived from a sparse row-normalized affinity matrix, and not the full covariance matrix.

For data that is a long-range experiment and not trial-based, it is possible to divide the data into *N* shorter subsets in time. The advantage of this is that short-term correlations are more accurate in defining similarity and less sensitive to noise.

## 3. ROI Extraction

We propose a novel clustering method based on the spectral embedding of the high-dimensional data-points. We present our method in the context of ROI extraction in calcium imaging data, where the clusters represent neuronal structures with distinct temporal activity, however this method is not tailored to a specific application. Our approach is motivated by spectral clustering, whose limitations we will briefly discuss.

The eigenvectors corresponding to the smallest eigenvalues of the normalized Laplacian

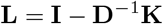

are employed in classical spectral clustering techniques, which propose selecting the first *k* eigenvectors for identifying *k* clusters, for example via k-means [18,29]. This is based on the property of the eigenvectors of the Laplacian to localize on clusters in the data. Various methods try to align these eigenvectors with the canonical coordinate system, i.e., the axis {*e_i_*} [19, 22]. It has recently been established that for well-defined clusters, the first *k* eigenvectors compose an orthogonal-cone-structure, where each cone comprises a cluster [21, 22]. Note that these eigenvectors are also eigenvectors of a Markov matrix on the data defined as in (2), motivating our approach.

Yet, for data which varies in density and cluster size, typically the first *k* eigenvectors will not identify *k* clusters. Instead several eigenvectors may “repeat” on the same cluster [20, 30]. An additional limitation of traditional approaches is that they assume all points belong to a cluster, whereas in calcium imaging (and other biomedical imaging applications), there are pixels belonging to the clutter (background), which are not of interest, and whose proportion out of the full image plane can vary (see Fig. 1 and 2(a)). Applying k-means to the eigenvectors, for example, will result in these background points being randomly assigned to one of the clusters, as the clutter will be randomly spread on the unit *k*-dimensional sphere and clustered along with the objects of interest.

**Figure 1.**
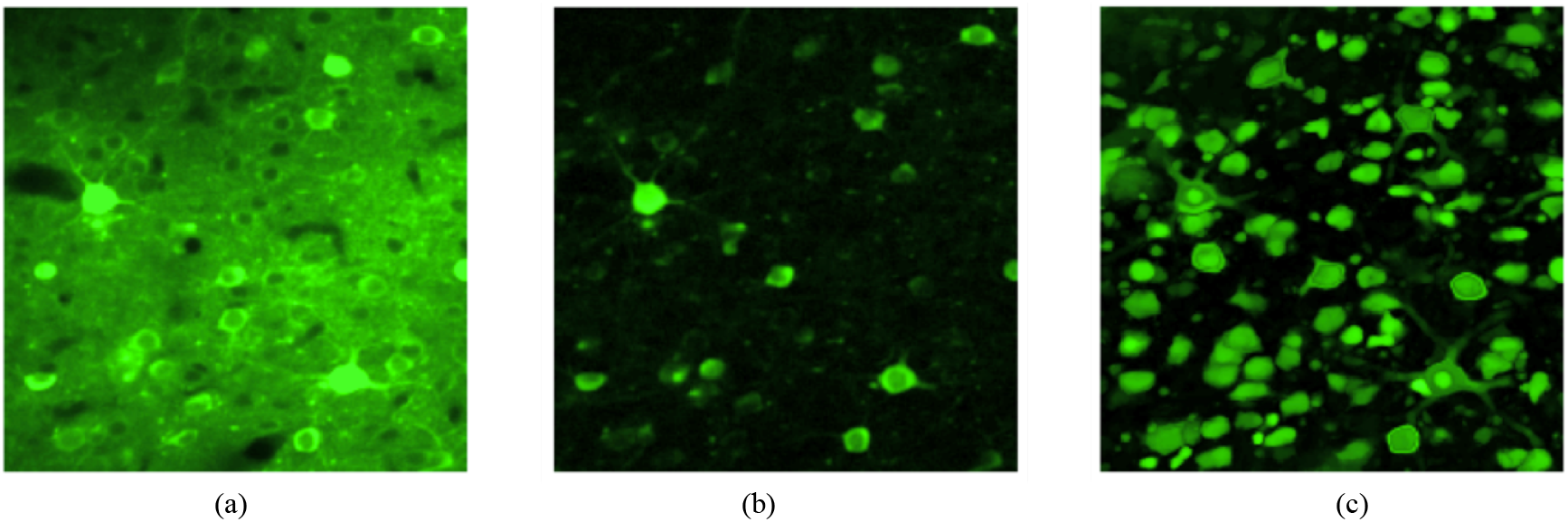
Figure 1. (a) Temporal mean image. (b) Temporal correlation image. (c) Embedding norm image. The clusters visualized in the embedding norm image correspond to the typical “donut” structure of neurons in the temporal mean image. Dendrites connecting to the soma are also seen with very high resolution, whereas they are barely visible in the other two images.

**Figure 2.**
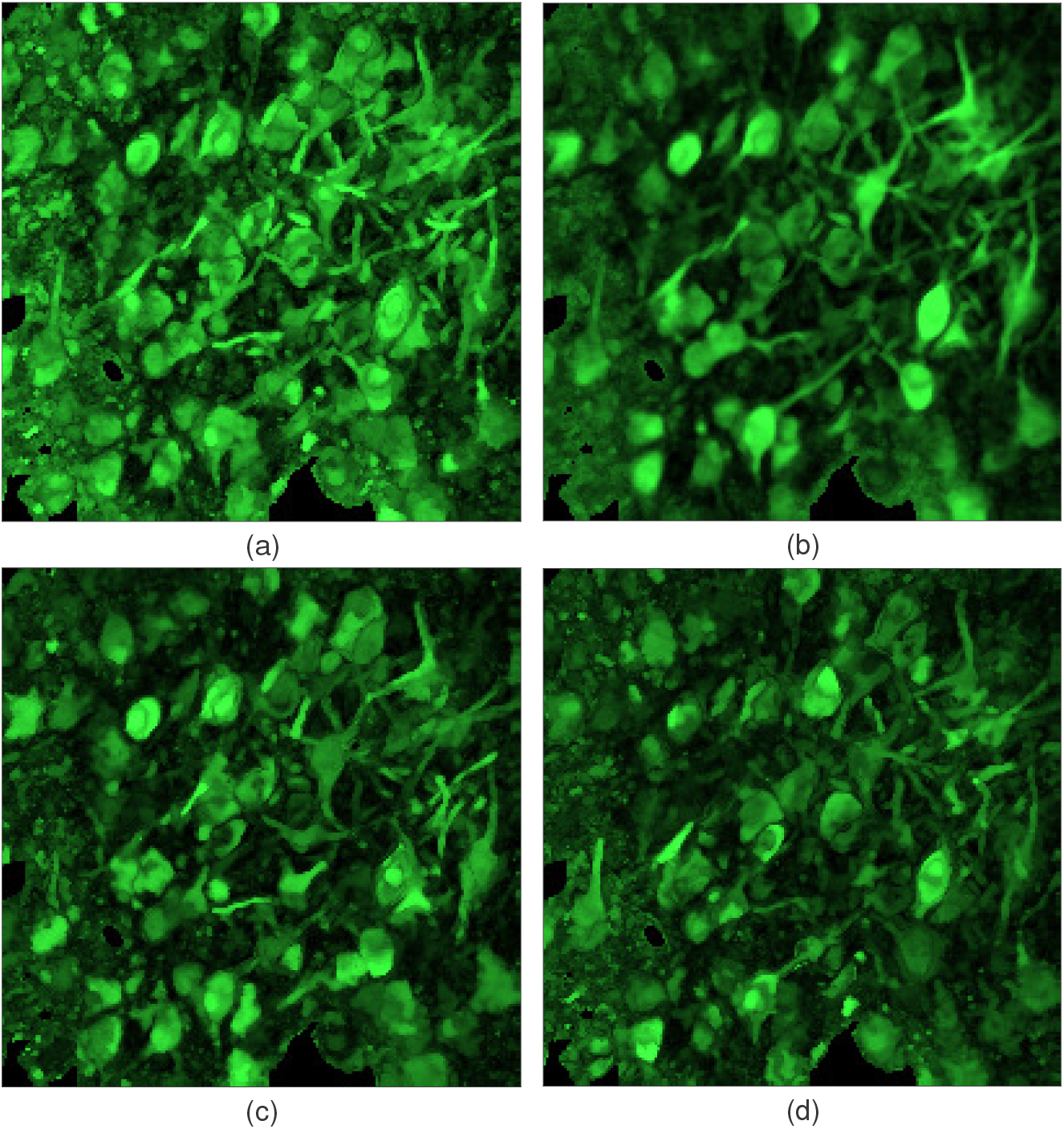
(a) Maximum embedding norm image, constructed by aggregating 20 trials where 50 eigenvectors have been calculated for each trial. (b) “Supervised” embedding norm image, where the embedding norm image has been calculated using a subset of eigenvectors from all 20 trials which are associated with ROIs we detected automatically. This enhances structures corresponding to the identified ROIs and removes interference form the background. (c)-(d) An unsupervised “random projection” construction of the embedding norm. Instead of using all 50 eigenvectors for each trial, we randomly select 10 out of 50 eigenvectors and calculate the maximum embedding norm of this subset. Both images are a random realization of this construction, where comparing the two reveals different structures. Some regions are enhanced while others disappear entirely. This enables a new multi-view visualization of data.

In Fig. 3(a) we plot the first three eigenvectors of **P**_1_: *ψ*_1_*, ψ*_2_*, ψ*_3_, for the first trial in the dataset corresponding to Fig. 2. Each point in the embedding space corresponds to a pixel in the image plane. This demonstrates the nature of the eigenvectors to localize on a cluster or subset of clusters in the data, while tending to zero on the rest of the data. The eigenvectors form three distinct branches, where each such branch corresponds to a small subset of clusters in the data, visualized in Fig. 3(b). For each of the three branches, we map a set of points along the branch, labeled in black, back into the image plane, revealing either single or multiple neurons in each branch. Meanwhile, the majority of points lies at the origin. Note that these branches mix different neurons together.

**Figure 3.**
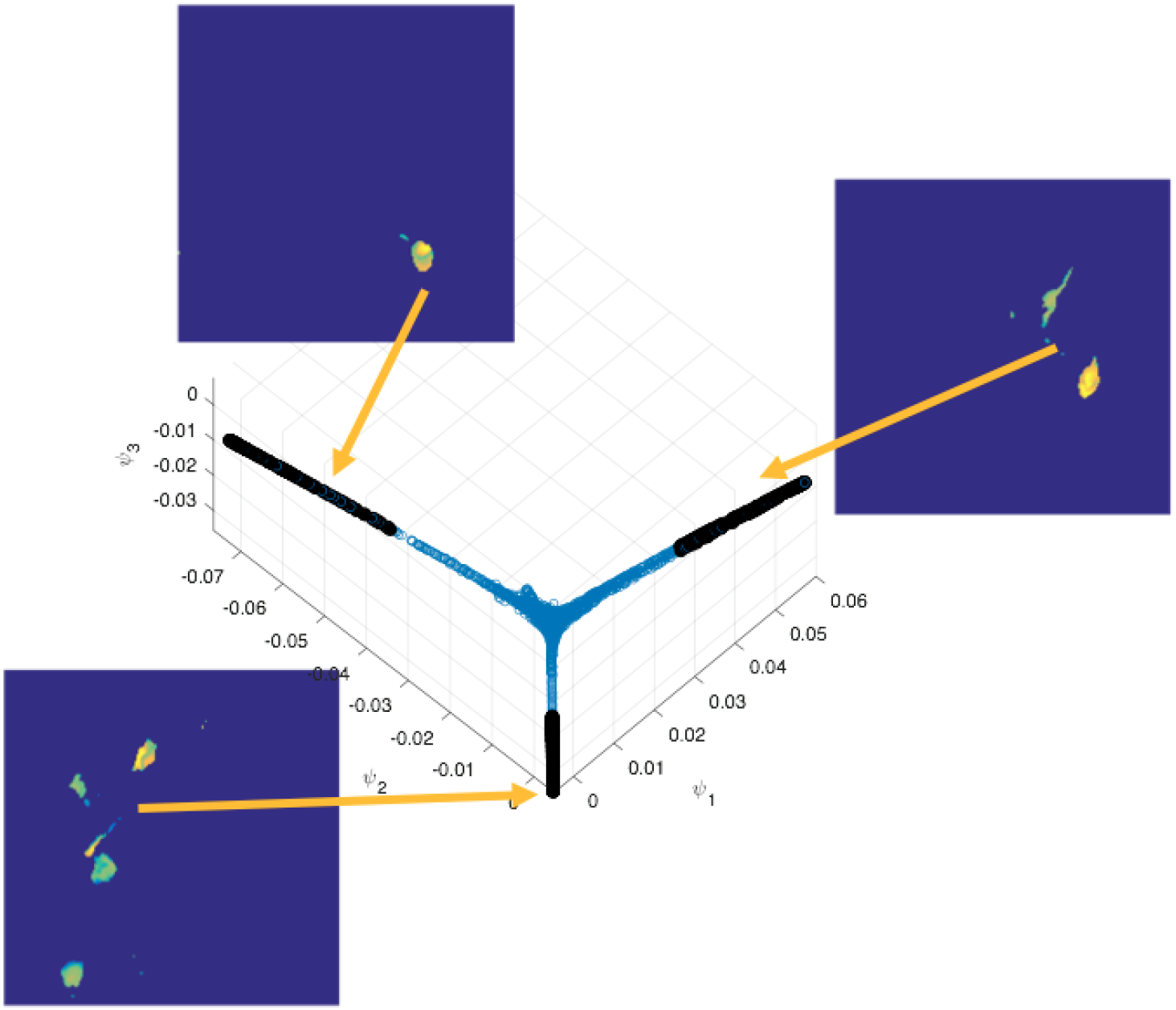
First three eigenvectors of a trial corresponding to the dataset in FIg. 2. Each point corresponds to a pixel in the image plane. The black points along each branch are visualized as a mask overlaid on the embedding norm image, revealing that each branch in the 3D embedding localizes on one or more neurons in the image plane.

We propose to take advantage of the eigenvectors localizing on different clusters in the data, but not limit the considered embedding space to only the first *k* eigenvectors, but rather look deeper into the spectrum. To this end, we present a novel spectral clustering approach, Local Selective Spectral Clustering, and apply it to ROI extraction in calcium imaging. We first present a global construction that visualizes the neuronal structures active in the video data as a 2D image. This image is used to initialize our clustering algorithm, as a measure for automatically detecting suspects belonging to clusters.

### 3.1. Embedding norm

An image summarizing the activity in a calcium imaging video is useful for both manually identifying ROIs, ground truth labeling, overlaying identified ROIs to determine what structure they capture and initializing ROI extraction methods as in [8]. Commonly used visualizations are the temporal mean image, temporal maximum image or a temporal correlation image [9] computed by

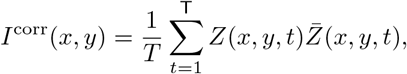

where *Z̄*_*n*_(*x, y, t*) is a spatial average of the pixels 4-adjacent (or 8-adjacent) neighbors (disregarding the image boundaries):

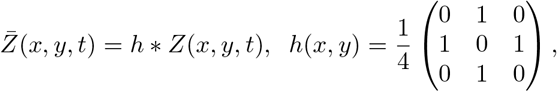

Where * indicates spatial convolution. Localized regions in the correlation image with high intensity correspond to strongly active cells, whereas localized regions with lower intensity correspond to neurons with lower intensity or other non-stationary processes. However, in complicated and noisy data this method yields a poor representation of the underlying structure.

We present a new 2D visualization based on the eigenvectors of the graph-Laplacian. As demonstrated in Fig. 3, clusters lie in branches or ideally in cones [21] in the eigen-space, with the clutter converging to the origin. Furthermore, recent results [31] indicate that a point *i* attaining the maximal magnitude of an eigenvector |*ψ*(*i*)| of the Laplacian operator on a graph lies far from the boundary of its cluster, i.e., close to the center of the cluster. Thus, visualizing the norm of the embedding as a 2D image serves both to suppress the background (since it has a small norm), while measuring how far each point is from its cluster boundary.

For pixel *i* in trial *n*, we calculate its norm in the embedding space, and view this as an image:

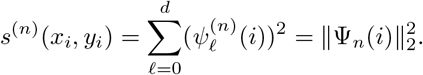

This image serves as a measure of how active each pixel is in a given trial *n*. In the case of multi-trial datasets, this measure is aggregated across all *N* trials, by calculating the maximum of 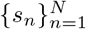:

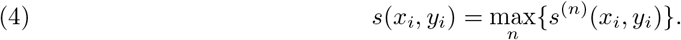

In this paper, we employ the embedding norm both globally and locally, on subsets of eigenvectors. The embedding norm has been considered as an object of interest for target detection in a supervised graph setting [32], and is closely related to the anomaly detection measure in [33].

In Fig. 1, we compare the temporal mean image (a), correlation image (b) and embedding norm (c) for a publicly available dataset from the Neurofinder Challenge website (http://neurofinder.codeneuro.org/). This dataset is 1000 seconds long imaged at 8Hz. The embedding norm visualizes multiple structures that are barely visible in the correlation image (if at all), and compared to the temporal mean image, it has removed all the background activity while visualizing both somas and dendrites with sharp morphology. Somas visible as “donuts” in the temporal mean image are visible in the embedding norm as filled in clusters.

Figure 2(a) presents the max embedding norm for a multi-trial dataset consisting of 20 consecutive trials, using the first 50 eigenvectors for each trial. This reveals both neurons and dendrites, with overlapping structures. The embedding norm can be used to reveal and enhance various structures in the data. In Fig. 2(b) we calculate a “supervised” embedding norm, where we do not sum over all eigenvectors, but rather only a subset of eigenvectors that are associated with an identified group of ROIs we extracted with our approach. This image enhances identifies structures, and removes the background interference. In a similar fashion we can consider, not a supervised construction but rather an unsupervised one. We randomly select 10 out of 50 eigenvectors for each trial composing the dataset and calculate the max embedding norm for this subset of eigenvectors. We present two such images in Fig. 2(c)-(d). Each image reveals certain structures, while others have vanished, or have reduced intensity compared to other regions in the image. Thus, as opposed to the correlation image which is “static”, here we can view the data from multiple viewpoints, each revealing new information and different levels of detail about the data. Viewing such random projections of the embedding is a novel construction (to the best of our knowledge) and its full implications and applications will be further explored in future work.

### 3.2. Local Selective Spectral Clustering

Having demonstrated the capability of the eigenvectors to visualize structure in the data, we present a new clustering approach for ROI extraction, inspired by spectral clustering [18, 29]. Our approach differs from traditional spectral clustering techniques in three aspects. First, instead of employing a global embedding of the data, we aim to find a local subset of eigenvectors, which best separates a cluster from the rest of the points. Second, instead of clustering all of the data at once, we present a greedy method that iterates over the different clusters. This allows for a stopping procedure so clustering the clutter can be avoided. Finally, we allow for multiple cluster membership, thereby detecting overlapping clusters. Since a point can be “expressed” in different subsets of localized eigenvectors, we can associate a point with more than one cluster. In calcium imaging, due to the axial resolution and the projection of a 3D volume onto a 2D plane, high measured intensity in a pixel can be traced to multiple neurons at the measured depth. For the sake of simplicity we begin with considering only one trial and neglect the index *n* in the following derivations.

We begin with composing a list *l* of the pixels in the image, ranked by a measure of their activity. Thus, we sort all pixels according to their maximum embedding norm *s*(*x, y*) (4) in descending order. Another possibility to construct *l* is to provide a user-interface with which to mark suspicious regions, for example, by providing the user with the maximum embedding norm, however this is outside the scope of this paper. Either ROI candidate initialization proposed in [8, 14] can also be applied to construct *l*. Since we iterate only over points with high embedding norm, we essentially ignore the clutter.

For a suspect point *i*, we find a subset of eigenvectors which form a “selective viewpoint” in which the suspect point and its corresponding cluster can be separated from the rest of the points. Starting with the point at the top of the list *l*, we sort the eigenvectors based on decreasing order of magnitude on point *i*:

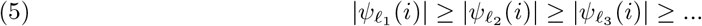

If the dataset is composed of multiple trials, we rank the eigenvectors from all trials together.

Next, we select only the first *d_i_* eigenvectors from this sequence, for example by thresholding the values with respect to |*ψ _𝓁_1__* (*i*)|, and set this subset to be *L_i_* = {𝓁_1_, 𝓁_2_*, …,* 𝓁*_d_i__*}. We denote the embedding of the data in this viewpoint as

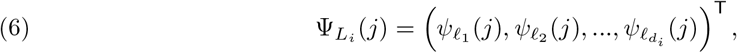

and term it a “selective viewpoint”. In this manner we find low-dimensional viewpoints in the embedding space, which best isolate a specific cluster. Points which belong to the same cluster as point *i*, will form a branch in the embedding space, where the maximal endpoint of this cluster is the centroid of the cluster, as indicated in [31]. Other clusters may occupy branches projecting into different directions in this space, while most points will collapse to the origin, due to the localization property of the eigenvectors. By taking into account embeddings from multiple trials, we leverage the redundancy of eigenvectors localizing on a cluster to better separate it from the background.

Having calculated *L_i_*, we want to select the points similar to **z***_i_*, identified by lying along the same branch as *Z_i_* in Ψ*_Li_*. We extract the points along the branch by comparing the distances of all points *Z_i_* to their distance to the origin in the selected subspace:

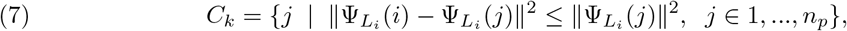

where *k* indexes the cluster, and *i* is the current suspect point. As most points lie at the origin, their distance to the origin, i.e., their norm, is smaller than their distance to the suspect point. Points closer to **z***_i_* are assigned to the cluster *C_k_*.

This is demonstrated in Fig. 4, for four different points and their selective viewpoint. Given a point we calculate its selective viewpoint, and then extract all points belonging to the branch that defines the point in this sub-space. In Fig. 4(a) we plot the first 3 leading eigenvectors of the selected viewpoint Ψ*_L_i__*. The black points indicate the extracted cluster and are mapped back into their corresponding pixels in the image plane in Fig. 4(b).

**Figure 4.**
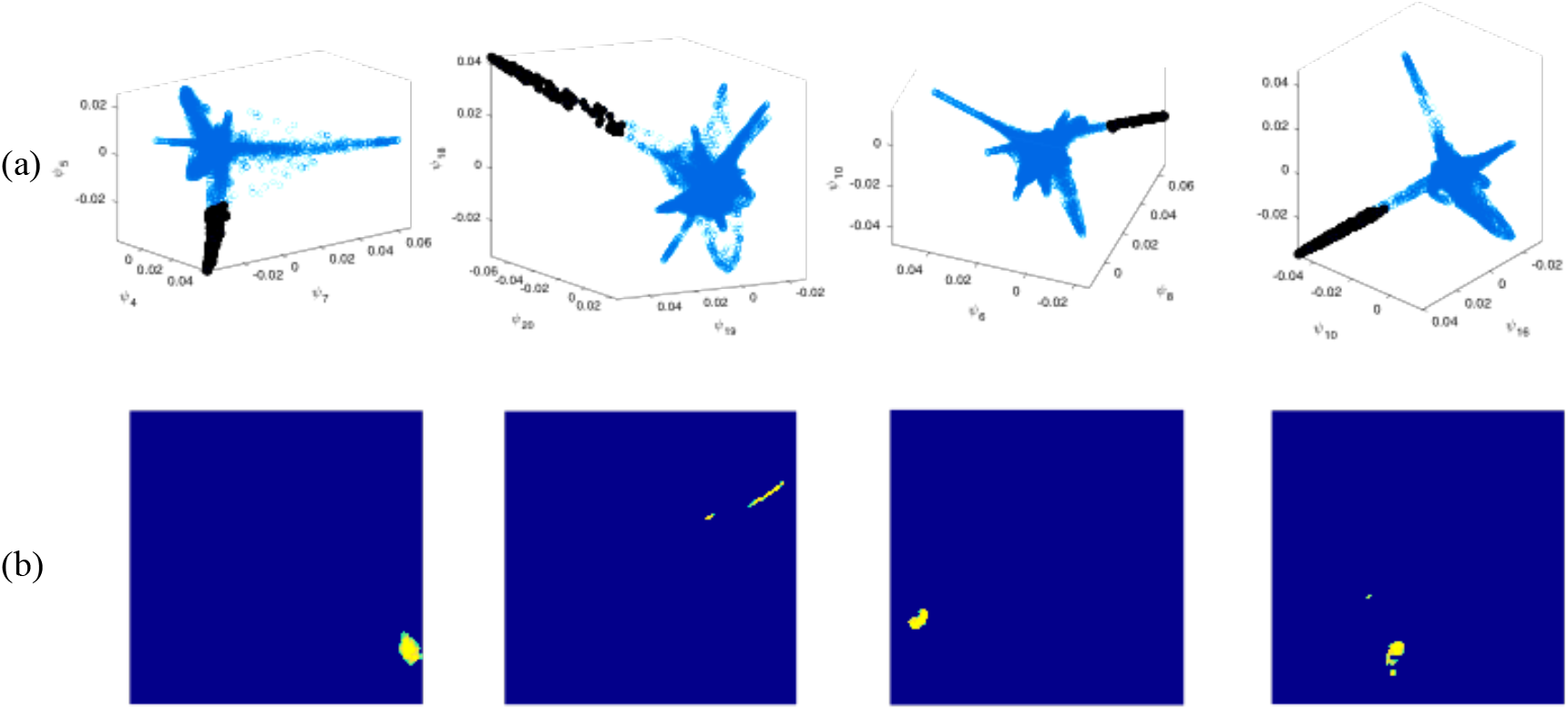
Local viewpoints extracting 4 different clusters. (a) The top three eigenvectors of the selective viewpoint Ψ*_L_i__* of a suspect *i*. Each local viewpoints contains several extending branches with the majority of points clustering at the origin. (b) The black points extracted from a single branch mapped back into the image plane, identifying either single neurons or a dendrite.

Next, we remove all points from *C_k_* that do not form a connected component with *Z_i_* in the image plane, since we assume that the ROIs are spatially compact. We then remove all points *j* ∈ *C_k_* from the list *l*, and apply the selective clustering to the point at the top of the list. Note that points already belonging to a cluster *C_k_* cannot initialize their own cluster, but they can belong to more than one cluster, thus enabling overlapping ROIs. We disregard a cluster *C_k_* if it is below a certain area threshold, i.e., we expect the ROIs to have a minimal spatial footprint. We stop once we have found *n*_clust_ clusters pre-specified by the user.

We next merge clusters based on spatial overlap *and* temporal correlation [14]. This procedure does not merge clusters that are spatially overlapping but are characterized by separate time-traces, i.e., overlapping neurons. The merging operation yields a final set of ROIs. We then rank the ROIs according to the product of the maximum values of their temporal and spatial components [14], so that the top-ranking ROIs have compact spatial support and strong temporal activity. Our clustering algorithm is provided in Alg. 1.

Our approach is related to the method in [34], where local “differential” viewpoints are constructed for anomaly detection. However, the viewpoints in [34] are created via random projections in a low-dimensional embedding space for the purpose of isolating single points from their neighborhoods to characterize anomalies, whereas we define the local viewpoint in a deterministic manner for the purpose of clustering, based on the properties of the embedding.

Note that the metric we use in (1) is based on temporal similarity of two pixels. One can adapt this metric to also incorporate spatial distances to suppress similarity between pixels that are correlated but spatially removed, however this requires adding a tradeoff parameter between spatial and temporal distances. In addition, using temporal similarity as in the current construction will mostly reveal active neurons. To detect non-active neurons, the metric can be further extended to incorporate spatial image features. Thus, the advantage of our clustering based approach is that different features and distances can be incorporated into the similarity kernel based on the structure of the data, and the required output clusters.

### 3.3. Cluster refinement

Once an initial cluster *C_k_* has been extracted, it can be further refined in the following way. We calculate a score for each eigenvector as

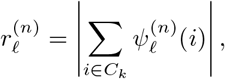

and rank the eigenvectors from all trials in descending value. Note that the absolute value is over the sum and not the individual values of the points. In this way, eigenvectors that oscillate on a certain cluster, thereby splitting it in the embedding space, are ranked lower than eigenvectors that have a smooth value across the cluster.

Selecting the top ranked eigenvectors defines a new viewpoint Ψ*_L_k__* for cluster *C_k_*. The centroid of the cluster is set to be the point whose embedding is the tip of the branch in this viewpoint, i.e., the point in *C_k_* with the highest embedding norm in Ψ*_L_k__*:

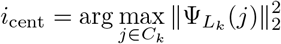

A new cluster is now extracted in this viewpoint by thresholding the distances of all points to *i*_cent_ in Ψ*_L_k__*. This refinement selects a viewpoint that is not defined by a single suspect point, but rather consistent across the initial cluster. Furthermore, the distances are not measured with respect to an arbitrary point in the cluster, but instead, to its centroid (defined as the tip of the branch).

In Fig. 5 we apply this refinement process to the initial clusters depicted in the top row, and obtain the clusters in the bottom row. This improves both somatic and dendritic ROIs.

**Figure 5.**
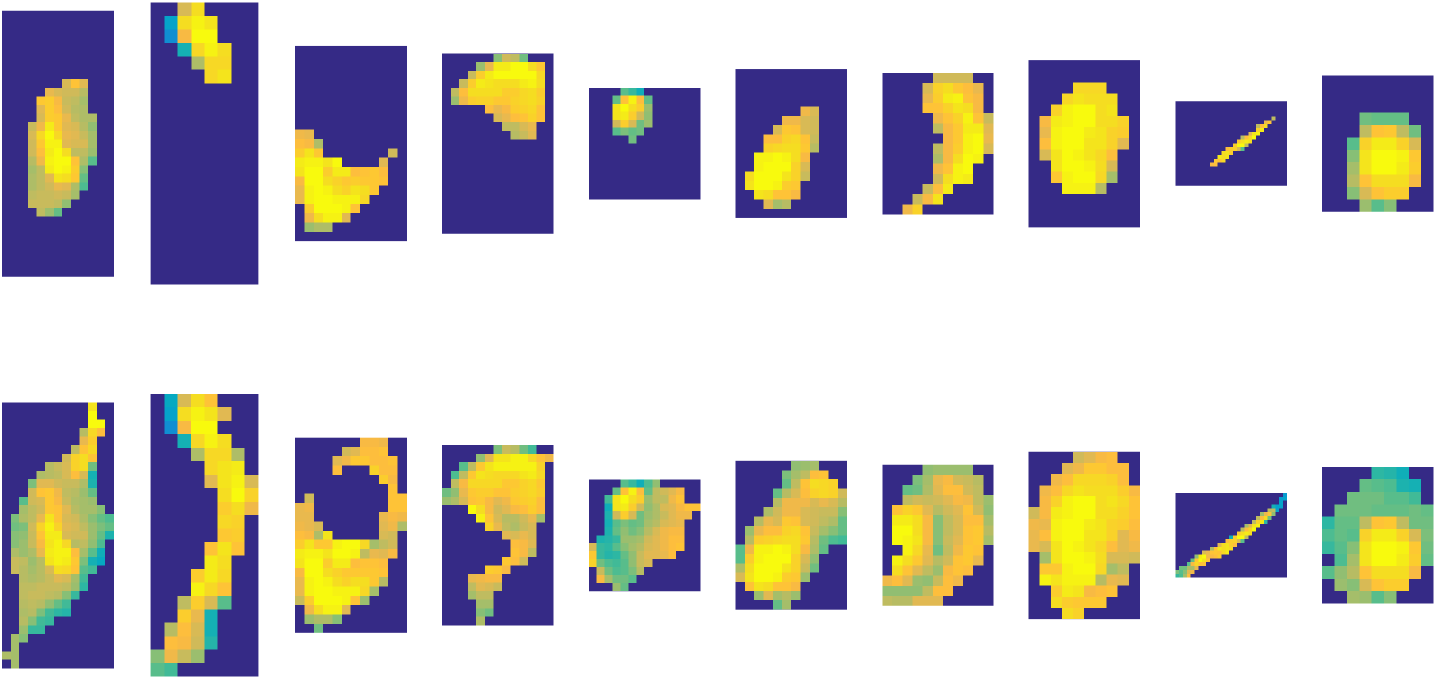
(top) Original extracted clusters from suspect points. (bottom) Refined clusters by calculating a consistent viewpoint and cluster centroid.

### 3.4. Large-scale data

In terms of practical considerations, constructing a graph for large-scale images with high-dimensionality has a high computational complexity. To address this, first we use approximate nearest-neighbor search [28] to construct a sparse affinity matrix in the spatial domain. Second, if the size of the image is too large, in order to speed-up and parallelize the processing, the image can be divided into sub-regions with overlap proportional to the size of a typical neuron. Since the neuronal structures are local in scale, each sub-region is analyzed separately. LSSC is applied to each sub-region in parallel and finally all clusters are grouped together and merged [14]. ROIs that have been split between two sub-regions, or found in both sub-regions in the overlapping area, will merge together due to their spatial overlap and temporal correlation. In addition, as in [15], since we do not have a non-negativity constraint [12, 14], we can preprocess the data to lower the dimensionality in the temporal domain by using a fast implementation of PCA [35].

In future work, we will explore an alternative construction for large scale datasets, by constructing a reference graph and applying out-of-sample-extension methods as in [32, 36, 37]. An initialization procedure can be used to select a reference set of suspicious points, and the embedding of all points in the image plane is dependent on their affinity to the reference set.

## 4. Temporal denoising

Our approach separates detecting spatial ROIs from calculating the temporal traces of each ROI, i.e., we do not solve a matrix factorization problem as in other methods. Thus, the temporal traces can be calculated based on the extracted spatial components using different methods. Here we present a greedy, local denoising scheme for calculating the temporal trace of each ROI. In addition, we leverage having calculated the spatial eigenvectors of the data to perform non-local diffusion filtering of the video data for enhanced viewing.

### 4.1. Local greedy temporal demixing and denoising

The time-traces of the ROIs suffer from two interferences. The first is the background activity due to the neuropil. The second is due to mixing in the signal when pixels belong to more than one ROI, i.e., a source separation problem. As opposed to a blind source separation problem, here we have knowledge of what pixels belong to more than one ROI and what pixels are unmixed.

#### Algorithm 1 Local Selective Spectral Clustering

~~~
**Input** Embedding of all pixels 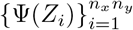, *s*(*x_i_, y_i_*), number of clusters *n_C_*
1: Create list *l* of all pixels sorted by decreasing value of *s*(*x_i_, y_i_*)
2: k = 0
3: **while** *k* < *n_C_* **do**
4:    Select *i* = arg max_*j*∈*l*_{*s*(*x_j_, y_j_*)}
5:    Select subset *L_i_* = {*ℓ*_1_, *ℓ*_2_, …, ℓ*_d_i__* } such that |*ψ _ℓ_*_1_ (*i*)| ≥ |*ψ _ℓ_*_2_ (*i*)*| ≥ |ψ _ℓ_*_3_ (*i*)| ≥ …
6:    *C_k_* = {*j* | ∥ Ψ*_L_i__* (*i*) *−* Ψ*_L_i__* (*j*)∥^2^ < ∥ Ψ*_L_i__* (*j*)∥^2^}
7:    Optional: refine *C_k_*
8:    *l* ← *l* \ *C_k_*
9:    **if** |*C_k_*| < *τ_area_* **then**
10:       discard *C_k_*
11:    **else**
12:    *k*← *k* + 1
13:    **end if**
14:    **end while**
15: Merge clusters based on spatial overlap and temporal correlation
**Output** Clusters {*C_k_*}
~~~

Given an ROI with no overlap, we apply a wavelet denoising scheme to denoise the temporal signal while maintaining the spiking profile of the activation potentials. We first calculate a weighted average of the ROI time-trace using all the time-traces of pixels belonging to the ROI

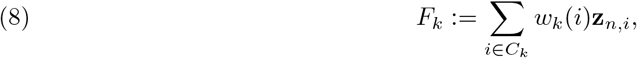

where we calculate the weights *w_k_*(*i*) based on the distance of point *i* form the cluster centroid *i*_cent_ in the local selective subspace Ψ*_L_k__* of cluster *C_k_*

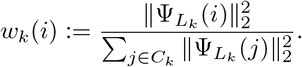

We then apply a wavelet denoising to *f_k_* using ‘db4’ wavelets, and hard-thresholding. We also apply spin-cycling for translation invariance, and denote the denoised time-trace by **f̂**_*k,n*_. For ROIs with overlaps, we develop a greedy PCA-based demixing algorithm. We view the time-traces of the pixels belonging to the ROI as different noisy realizations of the clean fluorescence signal of the neuron. Therefore, the first principal component captures the main temporal activity of the ROI. We sort the ROIs with ascending value of the amount of overlap. Beginning with a cluster with low overlap, we extract only its *n_p_* “pure” (unmixed) pixels. This defines a sub-matrix 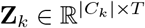, of the time-traces belonging to the pure pixels. We apply an SVD

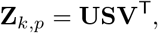

and calculate a projection matrix **P***_k_*= **vv**^T^ ∈ *ℝ^T^* ×*^T^*, where **v** is the first left singular vector in **V**. We then project all of the time-traces 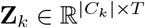 :

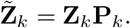

Thus, all the orthogonal components arising from the mixture with the other ROIs are suppressed. The time trace is calculated using weighted averaging (8) and wavelet denoising. We remove the mixture components from **Z** for all points belonging to *C_k_*:

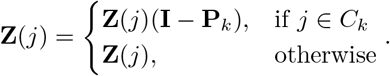

Therefore, when analyzing ROI *C_k’_* which overlaps with *C_k_*, we have removed the interference from the mixed pixels, so that only the signal associated with *C_k’_* remains.

In Fig. 6(a) we plot two overlapping ROIs, and in Fig. 6(b) we plot their time-traces, where the top plot corresponds to the left ROI (outlined in yellow) and the bottom plot corresponds to the right ROI (outlined in orange). The red plot is the average of the time-traces belonging to all pixels in the ROI. The left ROI has a higher intensity, so in the average trace of the right ROI, it is apparent that the few overlapping pixels greatly distort its time trace. Applying our demixing and denoising procedure yields the black time-traces, where the interference from the left ROI has been suppressed.

**Figure 6.**
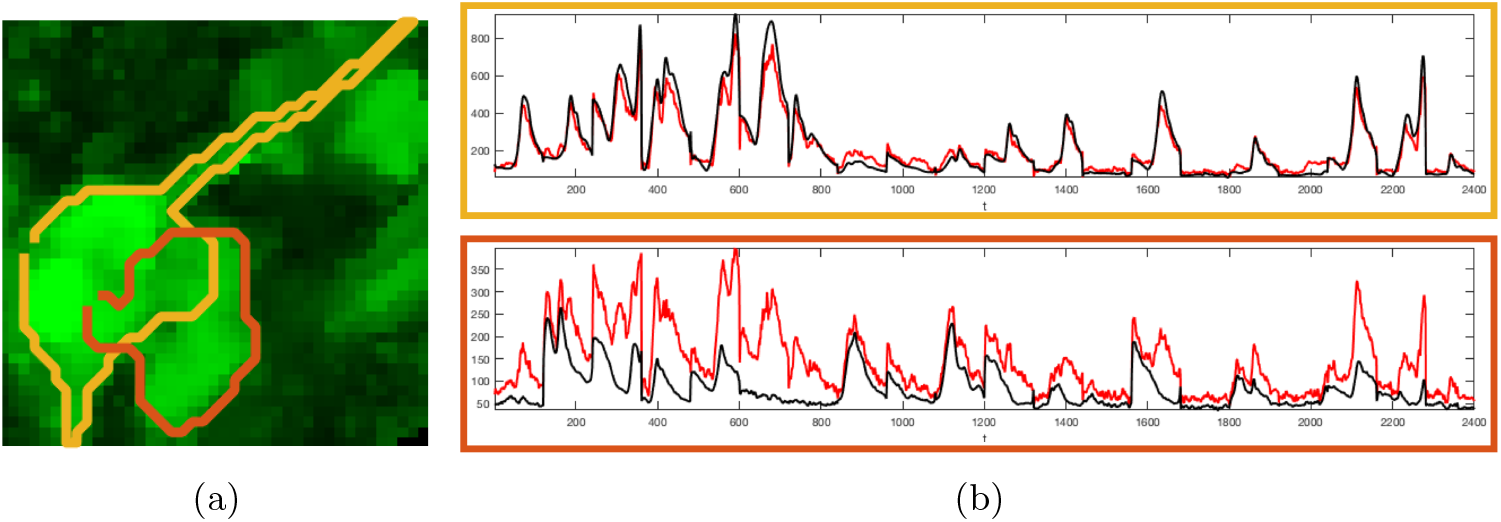
(a) Two overlapping ROIs overlaid on the embedding norm image. (b) The ROI time-traces, where the top plot corresponds to the left ROI (outlined in yellow) and the bottom plot corresponds to the right ROI (outlined in orange). We compare the average of all the time-traces belonging to the ROI (red) to the weighted average time-trace we extract after demixing and denoising (black). Applying the PCA demixing, interference from the left ROI has been suppressed in the overlapping pixels when calculating the time-trace of the right ROI. The wavelet denoising scheme yields smooth time-traces, with respect to the fixed-length trials of length 120 frames.

After the signal has been denoised, we calculate ∆*F*/*F*. The baseline florescence for *C_k_* in a single trial *n* is calculated using a subset of time frames *S_k_* corresponding to the florescent averages *F_k_*(*t*) with the 10% lowest values *F̄*_*k*_ = Σ_*t*∈_*_S_k__ F_k_*(*t*). Finally, the neuron measurement at each time frame is set as:

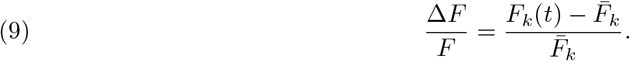

### 4.2. Global diffusion denoising

In addition to denoising the ROIs, we also propose a global denoising of the entire video volume using non-local diffusion filtering and employing the global spatial eigen-decomposition (3). Applying **P***_n_* (2) to the time-series **z***_n,i_* corresponds to applying a single denoising step by calculating the non-local mean of **z***_n,i_*[27]

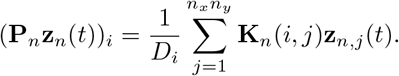

This amounts to a non-local *spatial* average of measurements **z**_*i*_[*t*] at time *t*. The denoising step can also be written as an eigen-basis expansion:

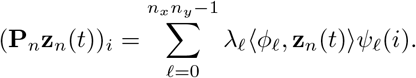

Due to the spectrum decay, we can discard the eigenvalues below a certain threshold and obtain an approximation by retaining only *d* eigenvalues and vectors:

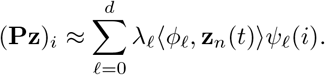

As with the embedding norm, since we discard the eigenvectors with low eigenvalues, which are typically related to noise, we suppress the noise in the images. Note that the global denoising we apply is a spatial denoising, where we leverage having calculated the eigen-decomposition of the spatial graph. Thus, we are averaging the measurements across different pixel locations at the same time-frame *t*.

In Fig. 7(a)-(b), we present images from several time-frames of the videos of two trials (after preprocessing of the dynamic range). Note the noisy nature of these images. In Fig. 7(c)-(d) we display the same time-frames after denoising, where the structure of neurons and dendrites has been preserved with sharp edges, while the background has been suppressed and smoothed out. The residual video **z – Pz** can be used to identify remaining neuronal activity not captured by the calculated eigenvectors, and to estimate the background signal.

**Figure 7.**
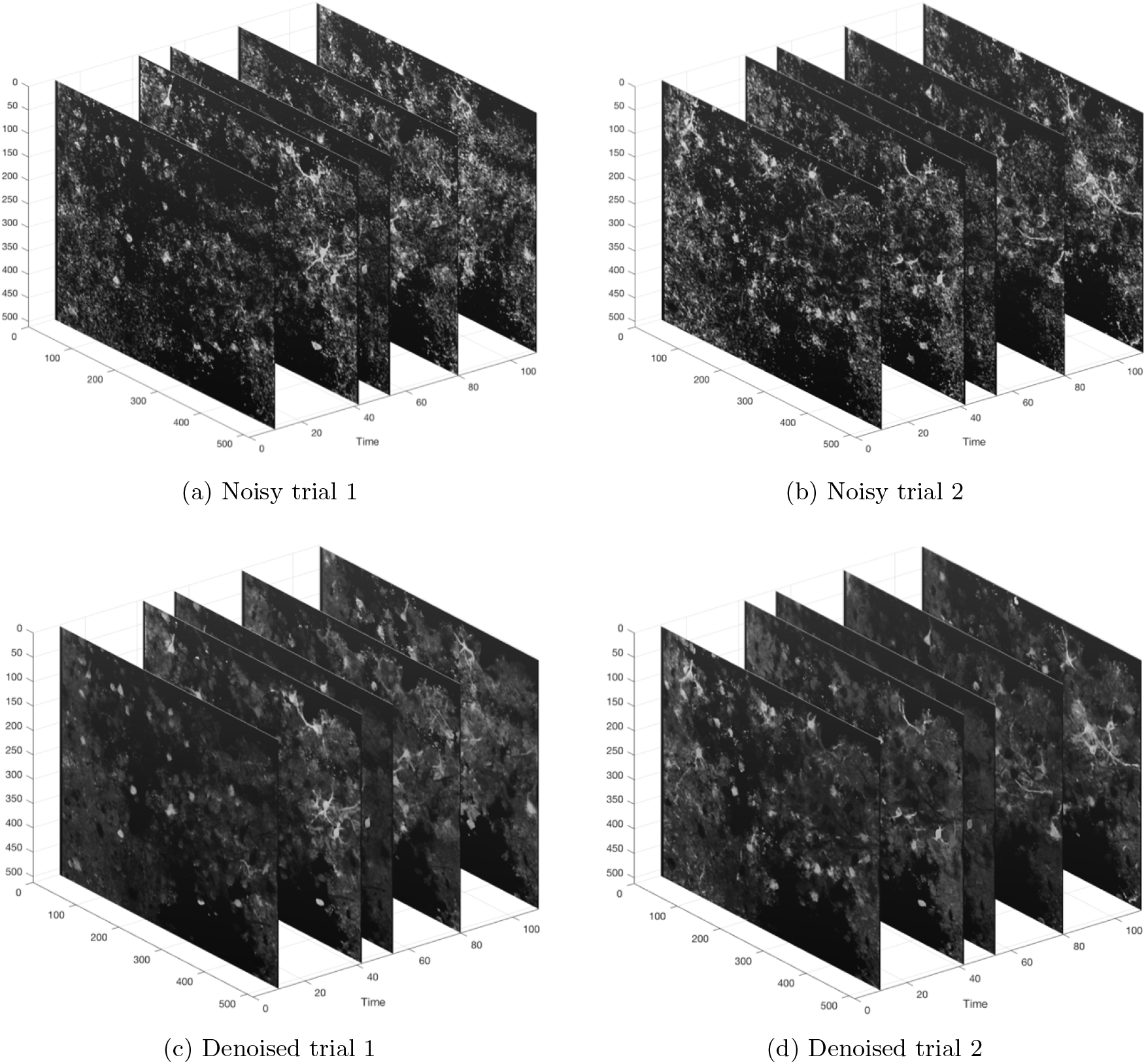
Spatial non-local diffusion denoising of two trials. (Top) Several frames from two trials (after preprocessing of the dynamic range). Then neuronal regions are highly contaminated by noise. (Bottom) The same frames after global diffusion denoising. The regions of interest remain as sharp bright structures while the noisy background has been suppressed.

## 5. Experimental Results

Our experimental data consists of repeated trials from a large population of primary motor cortex neurons from layers 2/3 and layer 5 acquired with two photon in-vivo calcium imaging.

### 5.1. Preprocessing

Before performing the spatial ROI extraction, we perform a preprocessing of the data to overcome the varying dynamic range across the image plane. We consider two possible preprocessing procedures: The first is normalizing the time-series of every trial **z***_n,i_* such that the lowest and highest 5 percent of values over times 0 ≤ *t* ≤ *T* are saturated, and the dynamic range is then mapped to [0, 1]. An alternative option is to z-score each time-series **z***_n,i_* by subtracting its mean and normalizing by its standard deviation (std). In either case, we then apply a 3 × 3 × 3 median filter to the volume, and flatten it into the matrix **Z***_n_*.

### 5.2. Somatic Imaging

We focus on neuronal measurements from the primary motor cortex (M1) acquired in a single day of experimental trials. The data is composed of *N* = 20 consecutive trials, where each trial lasted 12 seconds and is acquired at a frame rate of 10Hz, so that *T* = 120 for each trial. We analyze a region comprising 200 × 200 pixels.

The results are shown in Fig. 8. The number of clusters was set to *n*_clust_ = 200, and we calculated 50 eigenvectors for each trial, resulting in 1000 eigenvectors overall. We apply LSSC to extract the ROIs and after merging identified clusters and discarding those with area less than 50 pixels, 82 ROIs remained. The overall number of eigenvectors that were identified in different selective viewpoints is 693. The embedding norm using only these eigenvectors is displayed in Fig. 2(b).

**Figure 8.**
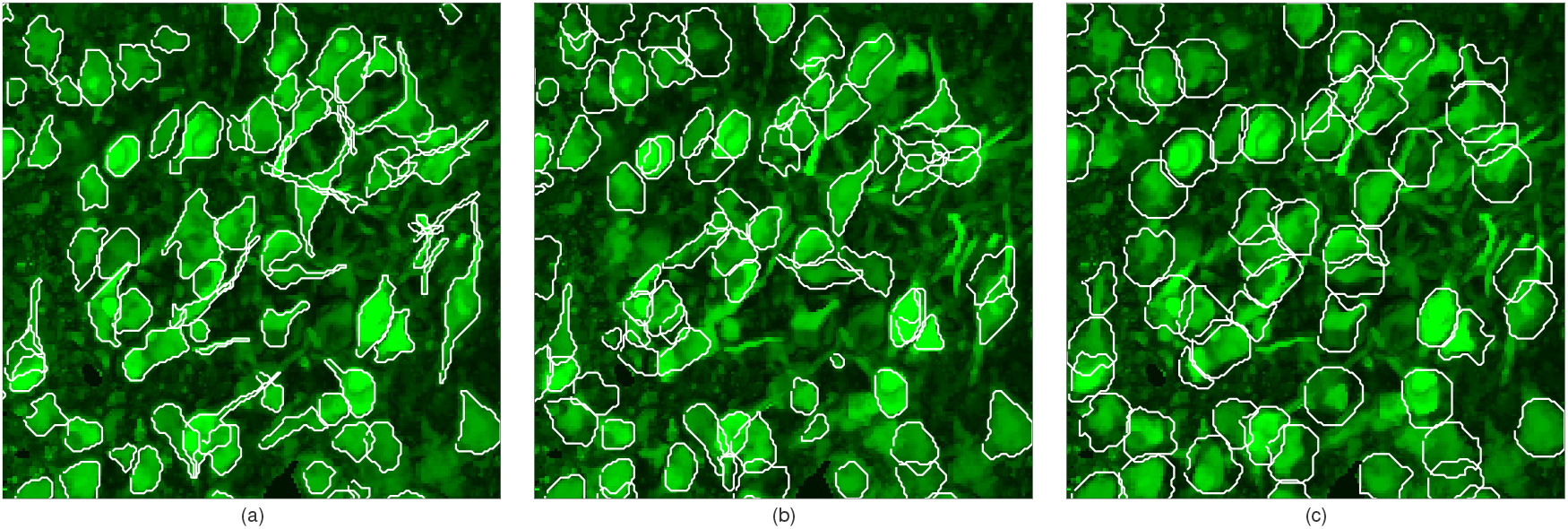
Analysis of 20 trials of length 12 seconds (10 Hz). The contours of the extracted ROIs are superimposed on the max embedding norm image, comparing LSSC approach (a) to CNMF [14] (b) and Suite2p [15] (c).

The ROI contours are depicted in Fig. 8(a), superimposed on the maximum embedding norm image. LSSC identifies neurons and dendrites with few visually-apparent false positives. The first 40 detected ROIs are shown in Fig. 9(a), superimposed on the max embedding norm image and demonstrating detailed morphological structures of both somas and dendrites. In Fig. 9(b) we plot the denoised time-traces of each ROI corresponding to Fig. 9a, where we reshape the time-traces as an image of size *N* × *T*. LSSC detects ROIs with various activity across trials, including ROIs with sparse support who rarely fire.

**Figure 9.**
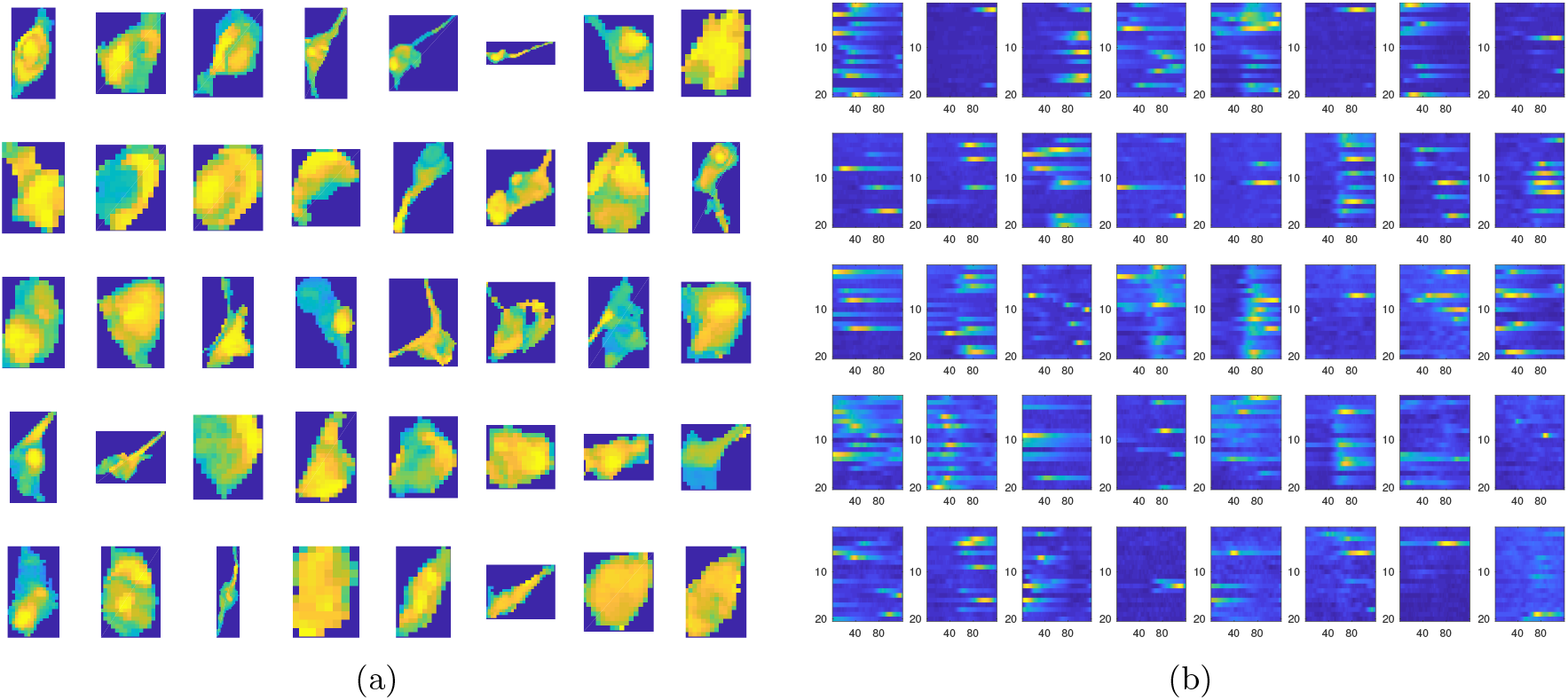
(a) First 40 ROIs extracted by LSSC. (b) Temporal traces reshaped as an image where each row is the temporal trace of a single trial consisting of 120 time frames (columns).

#### 5.2.1. Comparison

We compare LSSC to CNMF [14] in Fig. 8(b) and Suite2p [15] in Fig. 8(c). CNMF was initialized to detect 140 clusters and identified 86 ROIs after a post-processing procedure that includes merging and discarding components with a classifier. As both CNMF and LSSC require as input a number of clusters, both were initialized to output a similar number of ROIs to best identify differences between the methods in terms of what are the leading ROIs extracted by each method. For Suite2p, the number of ROIs is inferred by the algorithm and it detected 61 ROIs. We note that in our experiments both Suite2p and CNMF were sensitive to an input parameter each method requires regarding the expected size of the neurons, with the number and shape of output ROIs varying with slight changes of the parameter. LSSC on the other hand does not require an estimated neuron size for detection.

All the ROIs identified by Suite2p are identified by at least one of the other two methods, where the contours of the the ROIs detected by Suite2p tend to extend beyond those of the other two methods and capture more of the background. ROIs detected by LSSC have a finer contour adapting to the underlying structure. For CNMF, the high ranked ROIs have a fine morphology, but as indicated in [14], low ranked CNMF ROIs fail to converge to specific structures. Indeed, increasing the initial number of clusters in CNMF results in more and more overlapping ROIs and random ROIs with loose boundaries in the background. For LSSC, on the other hand, increasing the number of clusters is equivalent to examining structures with lower embedding norm, but the extraction of these structures is accurate. Note that the ROIs LSSC failed to detect all have low intensity in the embedding norm image.

Regarding identifying overlapping ROIs, CNMF had the best success rate, where in three cases it correctly identified overlapping ROIs that LSSC identified as a single ROI. Of these three cases, Suite2p failed to identify the overlapping ROIs in one of them. However, CNMF also tends to over-estimate overlapping ROIs, and in four cases detected at least 2 ROIs when there is only one. Regarding dendrites, LSSC displays capabilities of identifying fine dendritic structures in the data, even if they overlap other dendrites or soma. Suite2p detected three ROIs containing dendrites. In all of these cases, the dendrites extend from cell bodies that are present in the image plane and Suite2p split these into two separate ROIs, whereas LLSC identified them as a single ROI. As with the somas, these Suite2p ROIs extend beyond the dendritic structures into the background. CNMF did not detect individual dendrites.

#### 5.2.2. Large-scale images

Fig. 10 presents the analysis of a 512 × 512 of neuronal activity, across 20 fixed length trials lasting 12 seconds, acquired at 10 Hz. The video is split into 4 sub-regions in the image plane with horizontal and vertical overlaps, indicated by the dashed lines. Each sub-region (whose embedding norm is shown in images 1-4) is analyzed separately for ROI extraction, where around 100 ROIs where found in each sub-region, identified by the red contours. The ROIs are then merged in the full image (center) into 340 ROIs, and denoised. The sub-regions are overlapping in order to detect ROIs that may have been split between two or more regions, such as the examples indicated by the cyan and white ellipses. It should be noted that the image can be split into smaller sub-regions for the sake of improving run-time by constructing a smaller graph.

**Figure 10.**
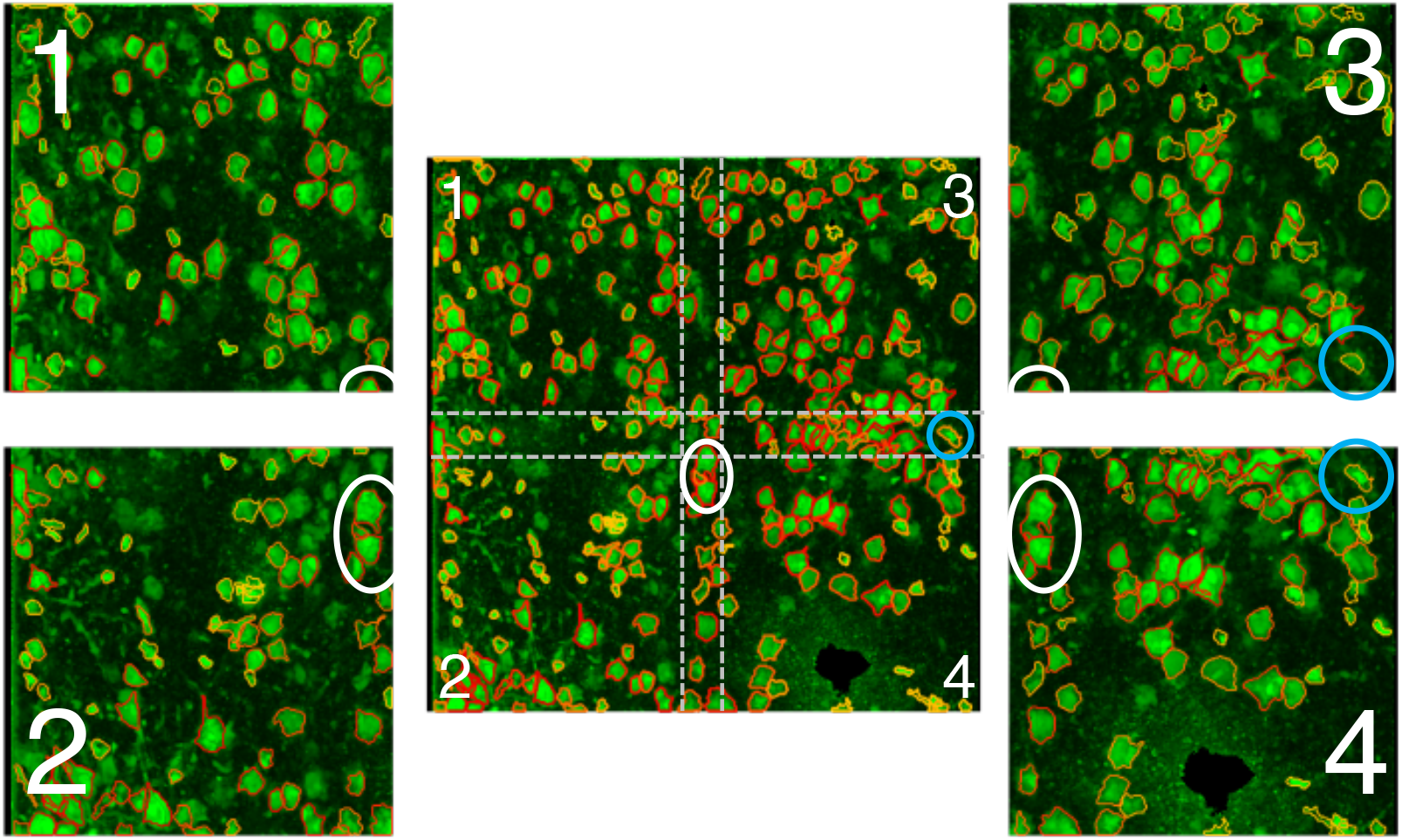
Analysis of large-scale images. The image is split into 4 sub-regions with horizontal and vertical overlaps (dashed lines). Each sub-region is analyzed separately (1-4) and the ROIs are then merged in the full image (center). An example of an ROI identified in sub-regions 3 and 4 is marked in cyan, and a group of ROIs identified fully in 2 and 4 and partially in 1 and 3 is marked in white.

### 5.3. Dendritic Imaging

We also analyze the extraction of apical dendrites. The data is composed of 35 consecutive trials, where each trial lasted 12 seconds. The image plane is 512a512 pixels acquired at a frame rate of 30Hz. In Fig. 11(a) we display the top 21 extracted ROIs, where each ROI is superimposed on an image of the temporal standard deviation. We plot the denoised temporal time-traces in Fig. 11(b), where we reshape the time-traces as an image of 35 trials × 360 times frames. Some dendrites share the same temporal structure; these sub-graphs can be automatically grouped and analyzed in a multi-scale organization method such as the one we presented in [23].

**Figure 11.**
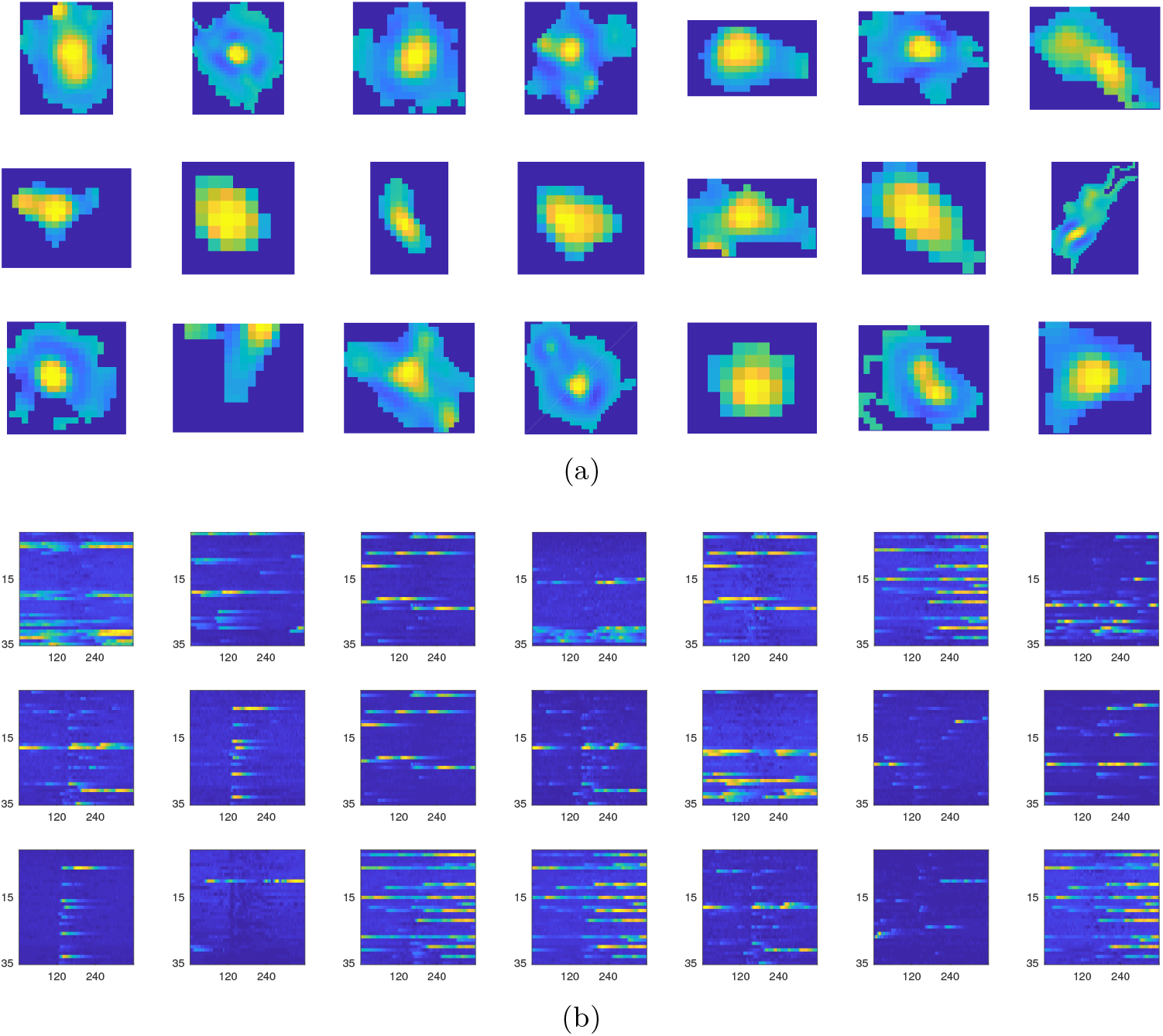
(a) First 21 ROIs. (b) Temporal traces reshaped as an image where each row is the temporal trace of a single trial consisting 360 time frames (columns).

In Fig. 12-13 we display two sub-regions of the dendritic image, with superimposed contours of the detected ROIs and their corresponding time-traces. The dendrites are partially overlapping, but detected as distinct ROIs. Again note the similarity between sub-groups of dendrites.

**Figure 12.**
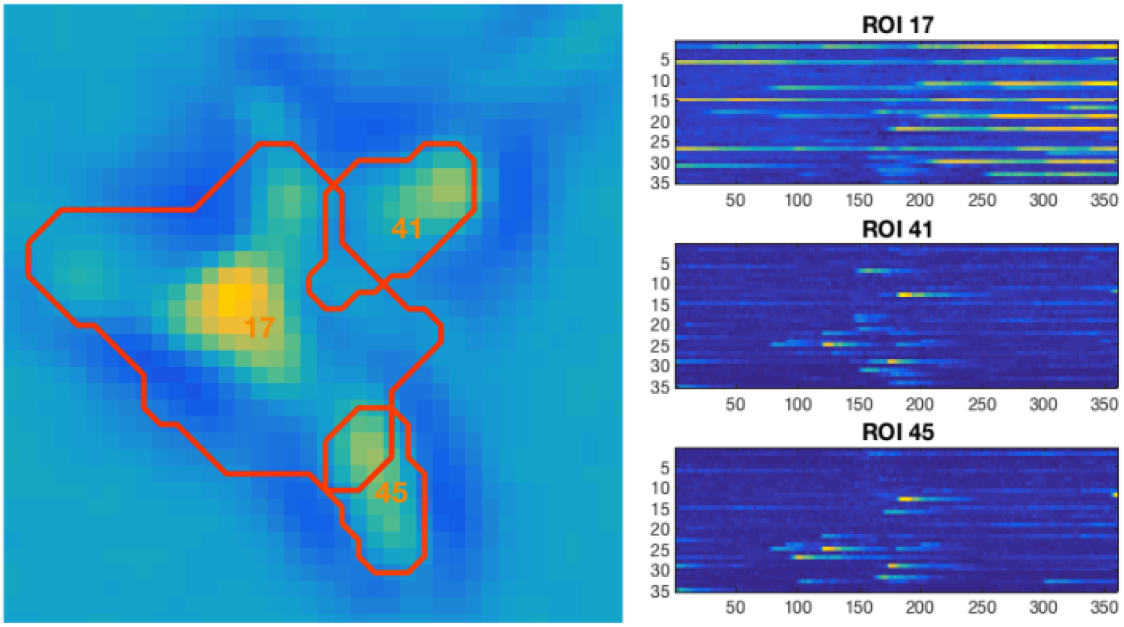
Sub-region of the dendritic image. Three partially overlapping dendrites (left) and their corresponding denoised and demixed time-traces (right).

**Figure 13.**
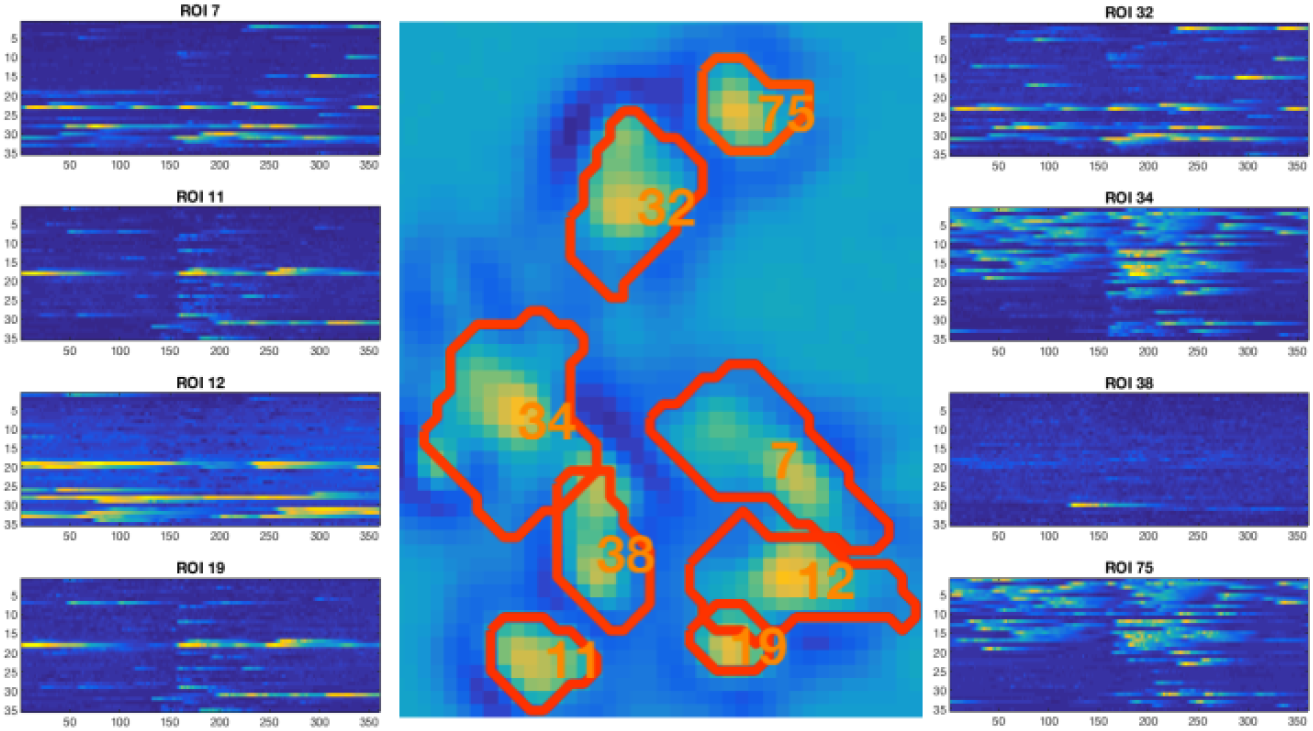
Sub-region of the dendritic image. Eight partially overlapping dendrites (center) and their corresponding denoised and demixed time-traces (left and right). Some of the dendrites share similar time-traces, which can be automatically grouped and analyzed in a multi-scale organization method such as [23].

## 6. Conclusions and future work

In this paper we presented local spectral methods for the processing of high-dimensional imaging datasets, and applied our framework to two photon calcium imaging data. We presented a new clustering method, Local Selective Spectral Clustering, inspired by classical spectral clustering, and capable of distinguishing overlapping clusters and disregarding clutter. We proposed using the low-dimensional embedding norm to visualize neuronal structure in calcium imaging videos. Finally, we developed a demixing and denoising scheme for the ROI temporal traces, employing wavelets and a greedy PCA-based source separation.

Two algorithmic choices we made that remain to be further formalized and explored are how to select the number of eigenvectors the selective viewpoint *L_i_*, and when to stop the greedy clustering process. Currently these are open questions allowing for flexibility in fulfilling the goals of ROI extraction. For the first question, we threshold the magnitude of the eigenvectors in comparison to the eigenvector with maximal value on the suspect point. One can also select a fixed number *L* of the large magnitude eigenvectors for each trial, yielding a selective viewpoint of size *LT*. Eigenvectors can be also selected based on mutual information, local linear regression, etc. [30]. Regarding the number of clusters, at present the user inputs a maximal number of clusters, however a stopping criteria can be introduced; for example, once the embedding norm *s*(*x_i_, y_i_*) of the remaining clusters is below a certain threshold, indicating the remaining points are clutter.

In this paper we have highlighted the low-dimensional point-wise embedding norm as an object of interest, briefly touching on possible supervised and unsupervised constructions of the norm that reveal various structures in the data. This will be further explored in a general high-dimensional data setting. We have also proposed a new clustering approach that can be applied to general datasets, beyond calcium imaging. We believe this approach has applications in other biomedical imagery, and even in other fields, such as demixing of substances in hyperspectral remote sensing imaging.

In the specific context of calcium imaging, one direction we intend to pursue in future work is extending our approach to the imaging of long-ranging dendrites, such as tuft dendrites. In this case the compactness of the support of the ROI and connectedness of all pixels no longer applies. A second direction is ROI extraction in one-photon data, which poses greater challenges due to high correlation of the neurons with the background.

## Acknowledgments

The authors thank Ronen Talmon, Xiuyuan Cheng, George Linderman and Nicholas Marshall for helpful discussions.

